# Evolutionary exploitation of PD-L1 expression in hormone receptor positive breast cancer

**DOI:** 10.1101/454447

**Authors:** Jeffrey West, Derek Park, Cathal Harmon, Drew Williamson, Peter Ashcroft, Davide Maestrini, Alexandra Ardaseva, Rafael Bravo, Prativa Sahoo, Hung Khong, Kimberly Luddy, Mark Robertson-Tessi

## Abstract

Based on clinical data from hormone positive breast cancer patients, we determined that there is a potential tradeoff between reducing tumor burden and altering metastatic potential when administering combination therapy of aromatase inhibitors and immune checkpoint inhibitors. While hormone-deprivation therapies serve to reduce tumor size in the neoadjuvant setting pre-surgery, they may induce tumors to change expression patterns towards a metastatic phenotype. We used mathematical modeling to explore how the timing of the therapies affects tumor burden and metastatic potential with an eye toward developing a dynamic prognostic score and reducing both tumor size and risk of metastasis.

## I. INTRODUCTION

Hormone-receptor positivity is observed in almost 75% of breast cancers, and these are frequently treated with aromatase inhibitors (AI) targeting the estrogen-receptor (ER^+^) pathway that fuels tumor growth in these patients. Here we examine neoadjuvant AI therapy prior to surgery and the impact of adding a concurrent immunotherapy. Although not traditionally considered a highly immunogenic cancer (1), ER^+^ patients that do not respond well to AI therapy have been shown to have a treatment-induced immune response, leading to the hypothesis that AI non-responders could benefit from combination immunotherapy. Programmed death ligand 1 (PD-L1) is a surface protein expressed on tumor cells and immune cells that acts to shut down the adaptive T-cell response to a tumor, and checkpoint inhibitors (CI) can be used to reverse this immunosuppressive effect (2). At present, a first-of-its-kind clinical trial using combination AI and CI therapies in the neoadjuvant setting for ER^+^ breast cancer is accruing patients.

This study aims to generate hypotheses regarding the optimal scheduling of both drugs in order to minimize the ‘preoperative endocrine prognostic index’ (PEPI), which accounts for tumor size, cancer cell presence in the lymph, Ki67 expression levels, and hormone receptor status. The PEPI score is shown to be a prognostic indicator for relapse-free survival (RFS). Here, we introduce mathematical models which consider the essential elements of the PEPI score. In particular we focus on tumor size and metastatic risk status of the tumor. Patient data suggests that AI therapy increases the expression of both PD-L1 and chemokine receptor CCR7 in tumors (3), both of which are associated with increased metastatic risk (4–6). Analysis of the therapeutic interactions suggests that there may be tradeoffs between tumor size and metastatic potential, and to this end we have developed a suite of mathematical models to investigate this interaction with the goal of eventually developing a dynamic PEPI score.

Three questions of interest were considered and explored using mathematical modeling: 1) How does metastatic potential change with therapy and can the combination of AI and CI be optimized to reduce this, while still lowering tumor burden? 2) What mechanisms of induction for the expression of CCR7 are at play, and how do they change in space and time? 3) How will the PEPI score change with the addition of CI therapy to AI therapy?

## II. METHODS

In order to predict phenotypic evolution in response to therapy, two specific phenotypic traits were considered: CCR7 and PD-L1 expression. CCR7 expression is correlated to AI resistance while PD-L1 relates to the effectiveness of CI immunotherapy. The evolution of each phenotypic trait is modeled using evolutionary game theory, where competition parameters are stored in the payoff matrix A_j_ for each j^th^ drug (j=1 for AI; j=2 for CI). Each payoff matrix gives the benefit (i.e. “fitness”) between any two competing cell types (row versus column). The payoff matrix associated with phenotypic evolution of CCR7 under administration of AI is given by:

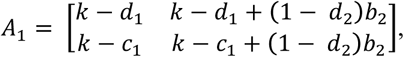

where *k* is immune kill rate by T cells, d_1_ is the kill rate by AI, c_1_ is the cost of developing resistance to AI, and b_2_ is the “cheater” benefit derived from being near a PD-L1 expressing cell, without self-expression of PD-L1. The first row and column represent the CCR7^−^ phenotype (i=1) and the second row represents the CCR7^+^ phenotype (i=2).

Similarly, the payoff matrix associated with phenotypic evolution of PD-L1 under CI administration is given by:

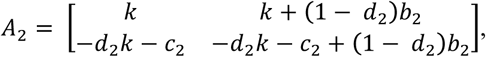

where c_2_ is cost of developing PD-L1 expression and d_2_ is the binary (d_2_ = 0 or 1) activation of T cells to target PD-L1 cells. The first row and column represent the PD-L1^−^ phenotype (i=1) and the second row represents the PD-L1^+^ phenotype (i=2).

Each j^th^ phenotypic trait (CCR7 or PD-L1) for each i^th^ cell type (negative or positive expression) can be tracked over time by a modified replicator dynamics equation:

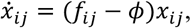

where cell type fitness is given by

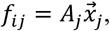

and average fitness is given by

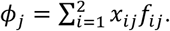

In addition to this game theory model, additional models were developed during the workshop to address our second and third questions of interest. The details of these models, while not presented here, are briefly referenced below.

## III. RESULTS

Figure 1 shows representative preliminary results for the three questions of interest, using varied mathematical modeling approaches. The hypotheses are discussed in more detail below.

**Figure 1:**
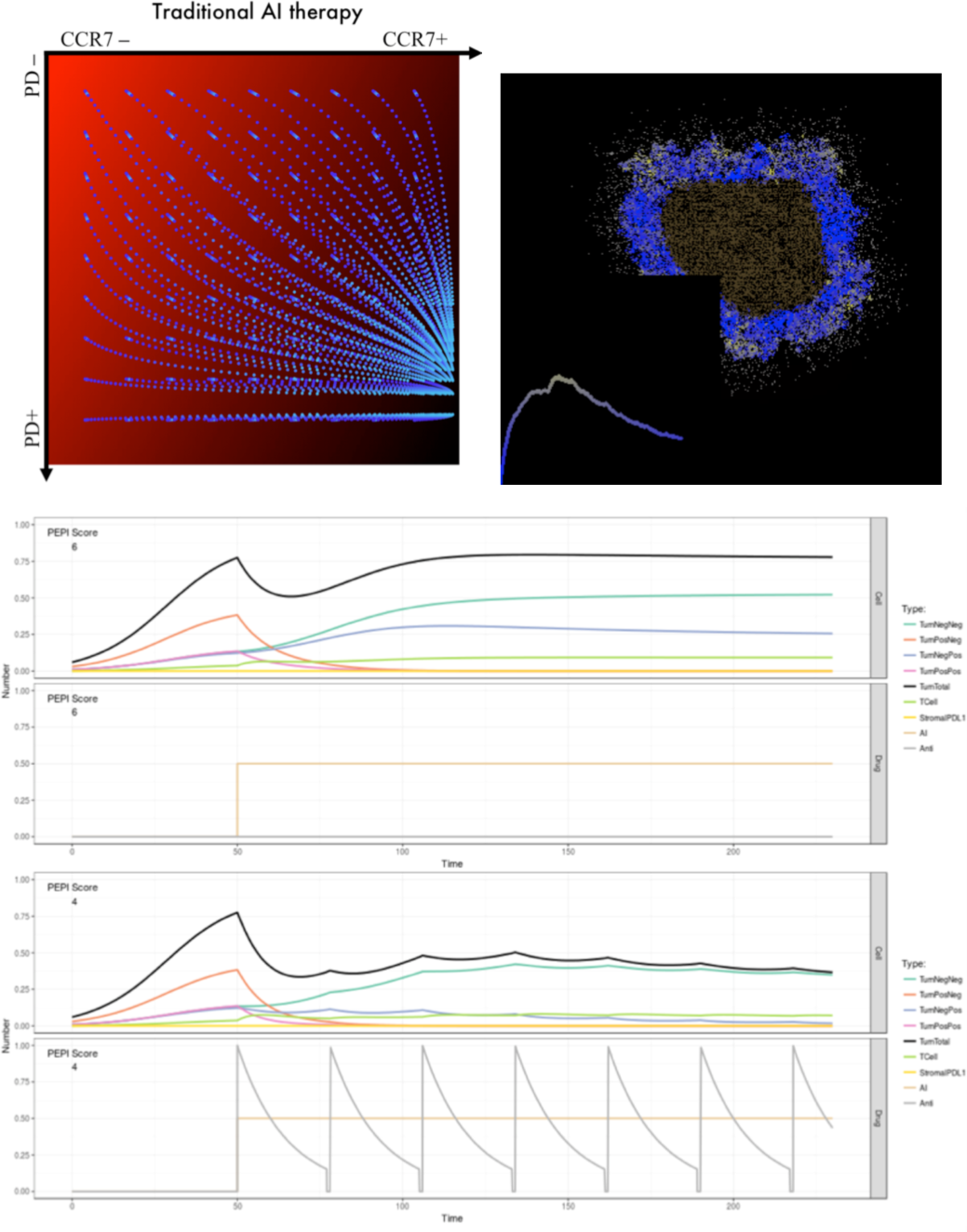
Representative results from the three models. Top left: A game-theoretic model uses trajectories (blue) through the expression space of CCR7 and PD-L1 under AI therapy to predict cumulative metastatic potential (red gradient). Top right: A cellular automaton model examines the possible mechanisms for inducing CCR7 expression, comparing spatial heterogeneity patterns with those seen in biopsies. Bottom: An ODE model predicts the improvement in PEPI score following combination AI and CI therapy.

### 1. Metastatic potential

The observation from patient data that CCR7 and PD-L1 expression are increased during application of neoadjuvant AI suggests that the metastatic potential of treated tumor cells may increase during the course of treatment. These markers are related to immune escape and cellular trafficking to lymph nodes and have been correlated with advanced stage disease. As described in the methods section, we use evolutionary game theory (7, 8) to investigate how the application of combination AI-CI therapy could be optimized to minimize this trade-off.

Since the time of the work presented here, which was generated during the week-long IMO Workshop, we have further elaborated and analyzed the model discussed in section II in a publication (9) that suggests that delaying the start of AI by one month can both decrease the tumor burden at surgery and reduce metastatic potential during the neoadjuvant period. Future work will include analysis of the associated clinical trial data looking at the response to this combination therapy.

### 2. CCR7 induction mechanism

Because the mechanism of increased CCR7 expression under AI therapy is unknown, we proposed to use a cellular automaton model to examine spatial heterogeneity of the expression pattern under different assumptions of the underlying mechanism, and compare these patterns to histology from patient samples. In the model, which was based on a hybrid automaton library (HAL, (10)), we simulated various mechanisms of CCR7 induction in response to application of AI therapy, including random expression, inflammation-driven expression, and estrogen-depletion-driven expression on ER^+^ cells. These different mechanisms produced different spatiotemporal patterns of expression pre- and post-therapy, which we can compare with histological samples from patients on the clinical trial.

### 3. PEPI score

The combination of AI and CI in ER^+^ breast cancer is novel and the trial at Moffitt is the first of its kind. To investigate how the addition of CI could improve the clinically-prognostic PEPI score, we built an ordinary differential equation (ODE) model that incorporates four tumor populations, T cells, and two therapies. The clinical PEPI score is based on tumor size, the expression of proliferation markers, involvement of lymph node metastases, and ER expression. In the ODE model, we measure these elements from the dynamics of the tumor subpopulations. Preliminary results suggest that addition of CI to the neoadjuvant therapy phase could improve PEPI scores by a value of 2 units, in the six month period before surgery. The model equations and an interactive simulation engine are available online at: https://ashcroftp.shinyapps.io/IMO-ODE-solver

## IV. CONCLUSIONS

Although immunotherapy is more well-known for its success as a monotherapy in melanoma and lung cancer, it is clear that there is a potential benefit to administering CI therapy to patients with ER^+^ breast cancer, particularly in combination with AI. In this workshop, we proposed that metastatic potential and tumor size, which are two key components of the clinical prognostic PEPI score, are both affected by the therapies in a way that sets up a tradeoff. Specifically, AI therapy will act to reduce tumor size but foment the expression of metastatic phenotypes. Addition of CI allows a second control to minimize the negative aspects of the trade-off. By using three modeling modalities, we can investigate schedule optimization to reduce metastatic potential, compare the spatiotemporal patterns of CCR7 expression under different assumptions, and predict the potential improvement in PEPI score that might arise in the upcoming clinical trial. Future directions include the development of a modeling system that would allow for dynamic PEPI scoring during the neoadjuvant phase of therapy, with the potential to adapt the treatments to patient data in real time.

## ACKNOWLEDGMENT

We would like to thank the IMO Chair, Dr. Alexander Anderson, for organizing the 7th Annual Moffitt IMO workshop: Stroma, where this project was conceived. We are also extremely grateful to the Moffitt Cancer Center and the Moffitt PSOC for supporting this workshop through the NCI U54CA193489 grant.

## REFERENCES

1 Ali HR, Glont SE, Blows FM, Provenzano E, Dawson SJ, Liu B, Hiller L, Dunn J, Poole CJ, Bowden S, Earl HM, Pharoah PD, Caldas C. PD-L1 protein expression in breast cancer is rare, enriched in basal-like tumours and associated with infiltrating lymphocytes. Ann Oncol. 2015;26(7):1488–93. doi: 10.1093/annonc/mdv192. PubMed PMID: 25897014.

2 Swart M, Verbrugge I, Beltman JB. Combination Approaches with Immune-Checkpoint Blockade in Cancer Therapy. Front Oncol. 2016;6:233. doi: 10.3389/fonc.2016.00233. PubMed PMID: 27847783; PMCID: PMC5088186.

3 Turnbull AK, Arthur LM, Renshaw L, Larionov AA, Kay C, Dunbier AK, Thomas JS, Dowsett M, Sims AH, Dixon JM. Accurate Prediction and Validation of Response to Endocrine Therapy in Breast Cancer. J Clin Oncol. 2015;33(20):2270–8. doi: 10.1200/JCO.2014.57.8963. PubMed PMID: 26033813.

4 Mazel M, Jacot W, Pantel K, Bartkowiak K, Topart D, Cayrefourcq L, Rossille D, Maudelonde T, Fest T, Alix-Panabieres C. Frequent expression of PD-L1 on circulating breast cancer cells. Mol Oncol. 2015;9(9):1773–82. doi: 10.1016/j.molonc.2015.05.009. PubMed PMID: 26093818; PMCID: PMC5528721.

5 Cunningham HD, Shannon LA, Calloway PA, Fassold BC, Dunwiddie I, Vielhauer G, Zhang M, Vines CM. Expression of the C-C chemokine receptor 7 mediates metastasis of breast cancer to the lymph nodes in mice. Transl Oncol. 2010;3(6):354–61. PubMed PMID: 21151474; PMCID: PMC3000460.

6 Xu B, Zhou M, Qiu W, Ye J, Feng Q. CCR7 mediates human breast cancer cell invasion, migration by inducing epithelial-mesenchymal transition and suppressing apoptosis through AKT pathway. Cancer Med. 2017;6(5):1062–71. doi: 10.1002/cam4.1039. PubMed PMID: 28378417; PMCID: PMC5430102.

7 Basanta D, Gatenby RA, Anderson AR. Exploiting evolution to treat drug resistance: combination therapy and the double bind. Mol Pharm. 2012;9(4):914–21. doi: 10.1021/mp200458e. PubMed PMID: 22369188; PMCID: PMC3325107.

8 Hummert S, Bohl K, Basanta D, Deutsch A, Werner S, Theissen G, Schroeter A, Schuster S. Evolutionary game theory: cells as players. Mol Biosyst. 2014;10(12):3044–65. doi: 10.1039/c3mb70602h. PubMed PMID: 25270362.

9 West J, Robertson-Tessi M, Luddy K, Park DS, Williamson DFK, Harmon C, Khong HT, Brown J, Anderson ARA. The immune checkpoint kick start: Optimization of neoadjuvant combination therapy using game theory. JCO Clin Cancer Inform (accepted); biorxiv. 2018. doi: 10.1101/349142.

10 Bravo RR, Robertson-Tessi M, Anderson ARA. Hybrid Automata Library. bioRxiv. 2018. doi: 10.1101/411538.

